# An orexin-sensitive subpopulation of layer 6 neurons regulates cortical excitability and anxiety behaviour

**DOI:** 10.1101/2024.07.25.605138

**Authors:** Fernando Messore, Rajeevan Narayanan Therpurakal, Jean-Philippe Dufour, Anna Hoerder-Suabedissen, Luiz Guidi, Kim Korrell, Marissa Mueller, Armin Lak, David Bannerman, Edward O Mann, Zoltán Molnár

## Abstract

Cortical layer 6 (L6) neurons uniquely respond to orexin – a neuropeptide influencing arousal and emotion. We show that Drd1a-Cre+ neurons in the prefrontal cortex are selectively sensitive to orexin and regulate prefrontal network activation in vitro and in vivo. Chronically silencing these neurons impairs orexin-induced prefrontal activation and reduces anxiety-like behaviour, indicating that orexin-responsive L6 neurons modulate emotional states and may be a substrate for anxiety regulation.

## Main Text

Neocortical layer 6 (L6) contains diverse types of pyramidal neurons that likely mediate distinct circuit functions, including cortico-cortical projection neurons and those making either feedforward or feedback projections to thalamus (Thomson, 2010). In sensorimotor neocortex, it has been shown that a Cre-driver mouse line targeting neurons expressing dopamine receptor 1a (Drd1a-Cre) selectively labels a subpopulation of these L6 projection neurons, which are concentrated close to the white matter, termed layer 6b (L6b), and send ascending projections to layer 1 and feedforward projections to higher-order thalamus (Bourassa et al., 1995; Clancy & Cauller, 1999; Hoerder-Suabedissen et al., 2018; Killackey & Sherman, 2003; Viswanathan et al., 2017; Zolnik et al., 2020). This subpopulation of neurons is uniquely sensitive to the wake-promoting neuropeptide, orexin, and their optogenetic activation is sufficient to trigger awake-like network states in anaesthetised neocortex *in vivo* (Zolnik et al., 2023), suggesting a role for neocortical Drd1a-Cre+ neurons in regulating cortical arousal. However, it remains unclear whether the anatomical and physiological properties of the Drd1a-Cre+ neurons are conserved across higher association areas, or what behavioural function this deep Drd1a-Cre+ neuronal subcircuit might serve. We focused on the medial prefrontal cortex (mPFC) activity to address these questions. We first explored the anatomical distribution of Drd1a-Cre+ neurons in mPFC of Drd1a-Cre:Ai14 mice, using DAPI staining to determine the layer borders, with TBR1 co-staining used to confirm L6 (Figure 1a) and complexin-3 (Cplx3) co-staining to identify L6b (Figure S1). We found that Drd1a-Cre+ neurons can be found almost exclusively in mPFC L6 (98.79%) (Figure 1b).

**Figure 1:**
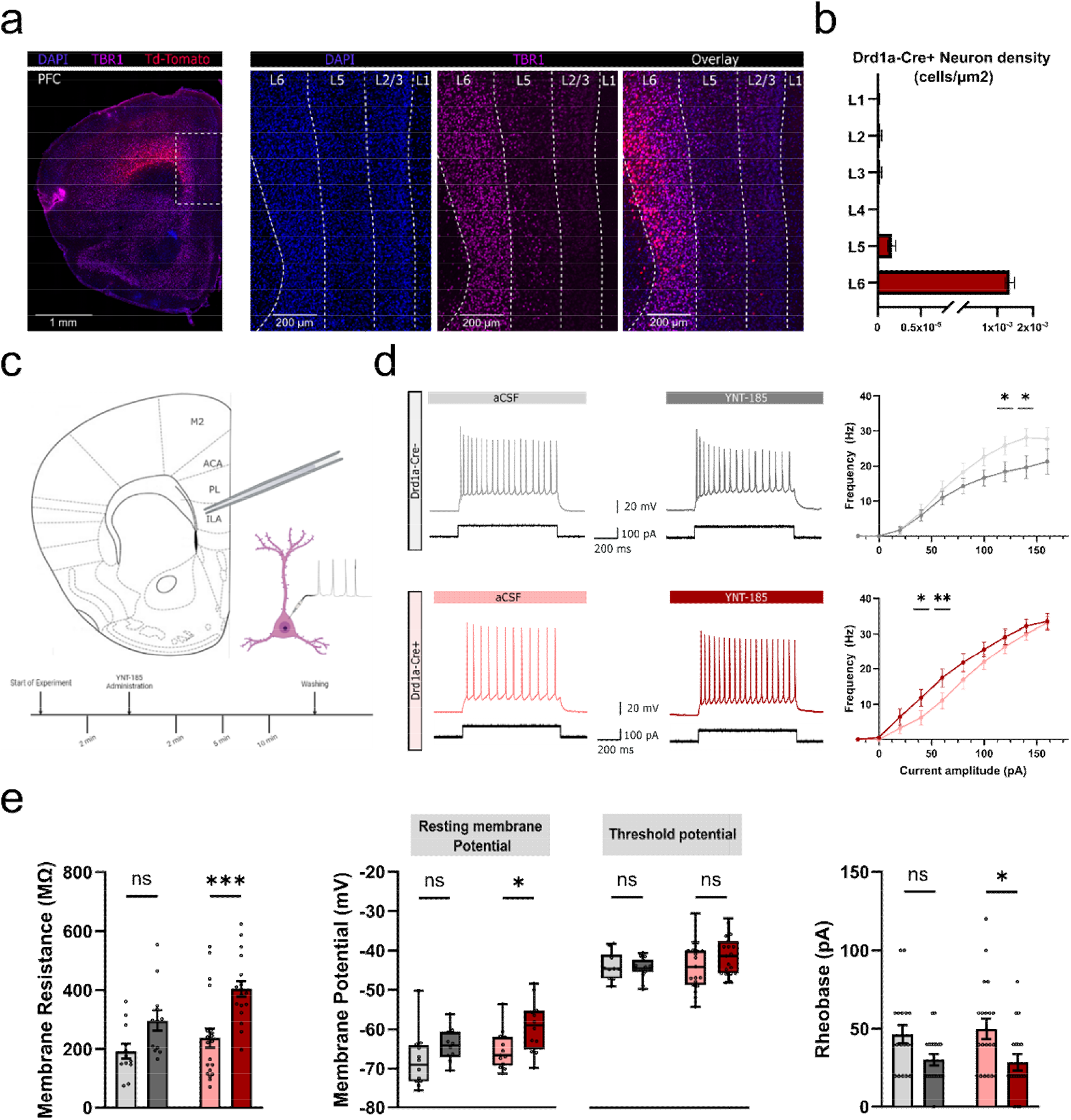
Drd1a-Cre+ neurons in layer 6 selectively respond to Orexin agonists in mPFC. **a)** Representative image of DAPI & TBR1 immunostained sections of Drd1a-Cre:Ai14 mice. These animals express Td-Tomato in the Drd1a-Cre+ neurons. Right panel shows a zoomed image of the medial prefrontal cortex (mPFC). DAPI staining was used to set the layer borders and TBR1 staining was used to delineate layer 6. **b)** Quantification of the number of cells per layer shows that Drd1a-Cre+ neurons are found almost exclusively in layer 6 of the mPFC. **c)** Schematic of the patch-clamp protocol. Drd1a-Cre+ neurons in the infralimbic cortex (ILA) were targeted for whole-cell current-clamp recordings and intrinsic properties were measured at baseline and at the 2, 5 and 10 minutes following application of 200 nM YNT-185. **d)** Example voltage recording after 100 pA stimulation in Drd1a-Cre-(gray) and Drd1a-Cre+ (red) neurons before and after the administration of YNT-185. Input-output curve shows the changes in spiking activity after increasing stimulation during baseline (aCSF) and administration of YNT-185. Drd1a-Cre-neurons (gray) become significantly inhibited at higher amplitudes (n = 12; p = 0.0199; Mixed effect analysis with Holm-Šídák’s multiple comparisons). Drd1a-Cre+ neurons (red) respond significantly higher at lower amplitudes after YNT-185 administration but plateau at higher amplitudes (n = 12; p = 0.0034; Mixed effect analysis with Holm-Šídák’s multiple comparisons). **e)** Membrane properties show significant changes after YNT-185 administration. Rheobase measurements show a reduction in the stimulus required to elicit an AP (n = 18; p = 0.0134; Two-way ANOVA with Šídák’s multiple comparisons correction). All numbers were reported as the mean with SEM. Error bars represent SEM, each datapoint is represented as a dot.

In order to explore the orexin-sensitivity of L6 mPFC neurons, we performed whole-cell current-clamp recordings from acute brain slices of Drd1a-Cre:Ai14 mice. In recordings from Drd1a-Cre+ L6 neurons, the application of the OX2R agonist, 200 nM YNT-185, increased intrinsic excitability due to an increase in membrane resistance and depolarisation of resting membrane potential (Figure 1c-e). These changes resulted in a reduced rheobase current, without a change in the voltage threshold for action potential generation (Figure S2). The increased spike rate for a given current step was associated with an increase in the half-width and a decrease in the spike amplitude, consistent with previous reports of frequency-dependent changes in action potential morphology (Geiger & Jonas, 2000; Kole et al., 2007; Kress & Mennerick, 2009; Shu et al., 2007) (Figure S3). Recordings from Drd1a-Cre-L6 neurons showed that YNT-185 only reduced the spike frequency at higher current steps (Figure 1c-g). Recent studies in the transcriptomic factors of layer 6 neurons have shown that subclasses of both excitatory and inhibitory L6 neurons can express orexin receptors (Henning et al., 2023), and thus the reduction in Drd1a-Cre-neuron firing rate could be due to the activation of L6 inhibitory neurons. While Drd1a-Cre+ L6 neurons may not be uniquely sensitive to orexin in mPFC, these results are consistent with a role for Drd1a-Cre+ L6 neurons in mediating orexin-induced increases in cortical excitability across sensorimotor cortex and mPFC.

While activation of OX2R increased the intrinsic excitability of Drd1a-Cre+ mPFC L6 neurons, it did not lead to sustained spiking activity, as observed with orexin application in the sensory cortex (Zolnik et al., 2023). To examine whether these changes in intrinsic excitability might be sufficient to activate cortical circuits, we performed 16-channel linear electrode recordings of the local field potential (LFP) in lower layers of mPFC of naïve mice under isoflurane anaesthesia (Figure 2a). Intraventricular injections of 1-2 µL 0.345 mM Orexin-B solution increased the burst frequency primarily at low-frequency power, both delta (0.5 - 4 Hz) and theta (4 - 8 Hz) frequency bands, but had no effect on higher frequencies (Wang & Mengoni, 2022) (Figure 2b). These effects were due to an increase in the occurrence of delta waves. This was also associated with an increase in the clustering of bursting events as has been seen for other neurotransmitters in the mPFC such as acetylcholine (Bloem et al., 2014) and dopamine (Laviolette et al., 2005; Lodge, 2011) (Figure S4). No changes in network activity were observed following intraventricular injections of 1-2 µL of saline solution (Figure 2a & b).

**Figure 2:**
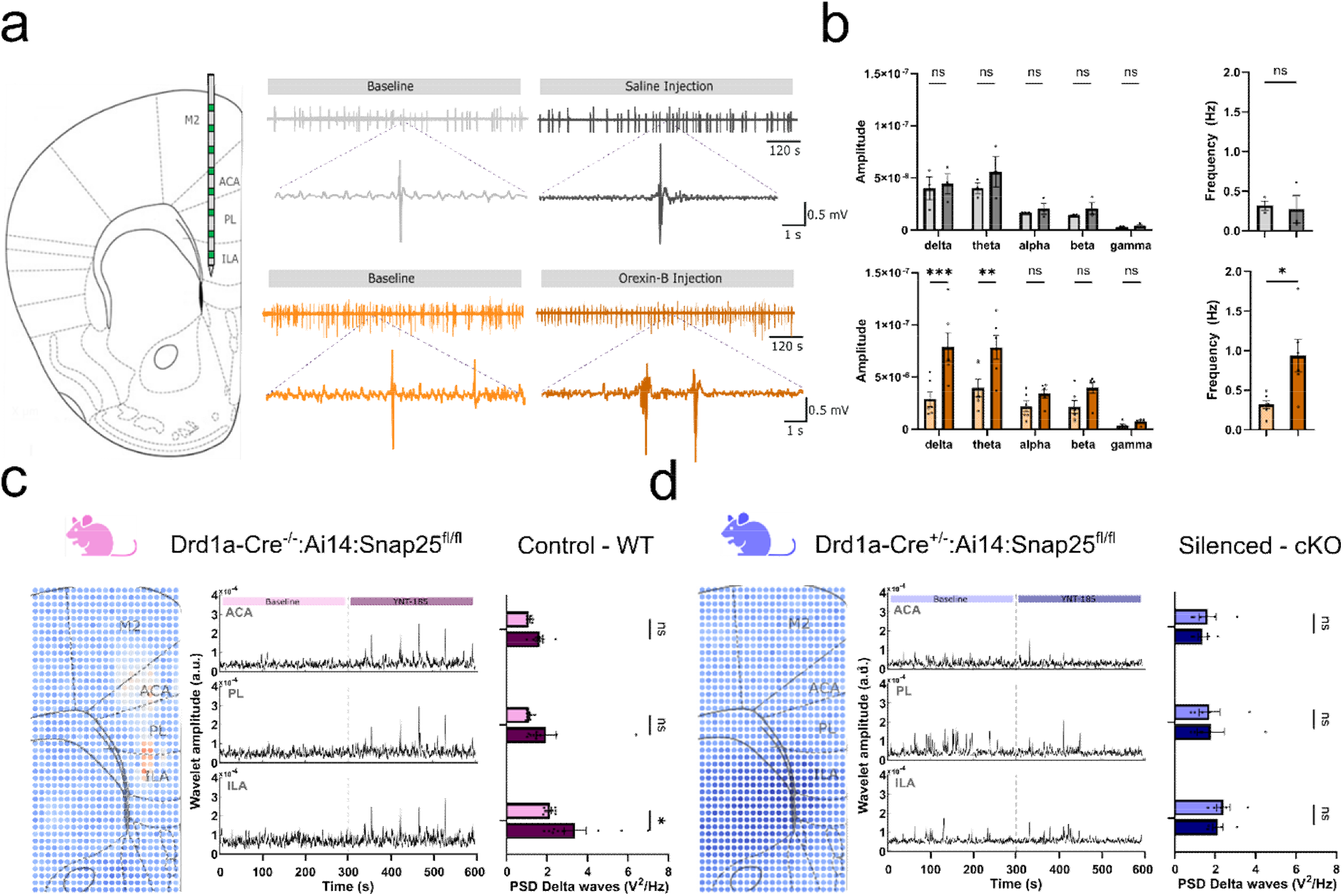
Chronic genetic silencing of Drd1a-Cre+ neurons abolish orexin response in mPFC. **a)** Schematic of the position of the linear electrode in the mPFC. This configuration allows for recordings in supplementary motor cortex (M2), anterior cingulate cortex (ACA), prelimbic cortex (PL), and infralimbic cortex (ILA). The spacing between the recording sites (green) is 110 µm. Example of electrode recordings in ILA through the experiment before and after the injection of saline (top) or orexin-B (boftom). **b)** Power spectral density (PSD) analysis of the recordings for the control (top) and the orexin experiments (boftom) in the ILA. No significant changes were observed for controls. Orexin administration significantly increased the spectral power measured in both delta and theta bands (n = 6; p = 0.0001 & p = 0.0024 respectively; Two-way ANOVA with Šídák’s multiple comparisons correction). This reflected an increase in burst frequency after the orexin injection compared to baseline and the saline injections (n = 6; p = 0.0308; Two-tailed Paired t-test). **c)** A PSD analysis of the network response to application of an ORX2 agonist, 200 nM YNT-815, in the control animals (Drd1a-Cre^-/-^:Ai14:Snap25^fl/fl^). Each dot represents an electrode in the array and warmer colours represent increases delta power after the drug administration. An increase in activity throughout mPFC can be seen. Wavelet transform analysis shows the change in delta power across time before and after YNT-185 administration. Right panel shows the changes in delta power per area. All areas show increased activity, which is significant in the ILA (n = 9; p = 0.0305; Mixed effect analysis with Šídák’s multiple comparisons). **d)** Same experiments were repeated in Drd1a-Cre-silenced animals (Drd1a-Cre^+/-^:Ai14:Snap25^fl/fl)^. There was a lack of YNT-185-driven cortical activation when these neurons were silenced. Wavelet transform analysis confirms the lack of increase in delta activity after the YNT-185 administration. All comparisons were done using mixed-effect Two-Way ANOVA analysis with Šidák correction All numbers were reported as the mean with SEM. Error bars represent SEM, each datapoint is represented as a dot.

To confirm that orexin-induced increases in delta activity were due to direct effects in mPFC, rather than indirect effects on subcortical arousal systems, we used high-density planar multielectrode arrays to record network bursting activity in acute slices of mPFC induced by application of 200 µM 4-AP. These bursts could be observed in the LFP, with power concentrated in the delta frequency range, and additional application of 200 nM YNT-185 increased both delta power and frequency of events (Figure 2c), as observed with orexinergic activation *in vivo*. To determine whether these effects were mediated by Drd1a-Cre+ L6 neurons, we repeated the experiments with mice in which these neurons had Snap25 ablated (Drd1a-Cre^+/-^:Ai14:Snap25^fl/fl^), thus eliminating action potential evoked release (Hoerder-Suabedissen et al., 2019). Functional silencing of Drd1a-Cre+ L6 neurons occluded the effects of YNT-185 (Figure 2d). This was not due to a simple decrease in the overall levels of synaptic activity, as YNT-185 increased activity throughout the mPFC in brain slices from mice in which layer 5 Rbp4-Cre+ neurons had been silenced (Figure S5). This suggests that Drd1a-Cre+ neurons selectively gate the orexinergic activation of mPFC.

The orexinergic activation of the mPFC has been associated with anxiety like behaviours (Cole et al., 2020; Karimi et al., 2019; Katzman & Katzman, 2022), such as an increased aversion to illuminated and exposed areas (Walf & Frye, 2007) as well as a reluctance to explore new environments (Bourin & Hascoet, 2003).

In order to explore whether Drd1a-Cre+ L6 neurons play a role in regulating anxiety, we compared the behaviour of mice with chronic silencing of Drd1a-Cre+ L6 neurons (Drd1a-Cre^+/-^:Ai14:Snap25^fl/fl^) with littermate controls (Drd1a-Cre^-/-^:Ai14:Snap25^fl/fl^) in two behavioural paradigms used to assess anxiety phenotypes: the Light-Dark Box (LDB) protocol and the Elevated Plus Maze (EPM) (Belzung & Griebel, 2001). In the LDB, animals tend to avoid the lit areas since they are a source of anxiety. Drd1a-Cre:silenced mice showed significant increases in the entries into the lit areas, as well as the total distance travelled in this area (Fig. 3a-b). Importantly, the distance travelled in the dark area remained similar across genotypes. Furthermore, assessment of spontaneous locomotor activity levels in novel photocell activity cages revealed no differences between the genotypes, demonstrating that the Drd1a-Cre:silenced mice were not hyperactive per se (Fig. 3c). Like the previous protocol, in the EPM, animals will prefer to remain in the enclosed arms instead of the exposed open arms (Fig. 3d). Consistent with a reduced anxiety phenotype, cKO animals showed a reduction in immobility time and an increase in the number of entries to these open arms (Fig 3e). Furthermore, these animals showed an increase in the distance moved and the time spent in the open arms, while the total amount of movement remained similar to the control animals (Fig. 3f).

**Figure 3:**
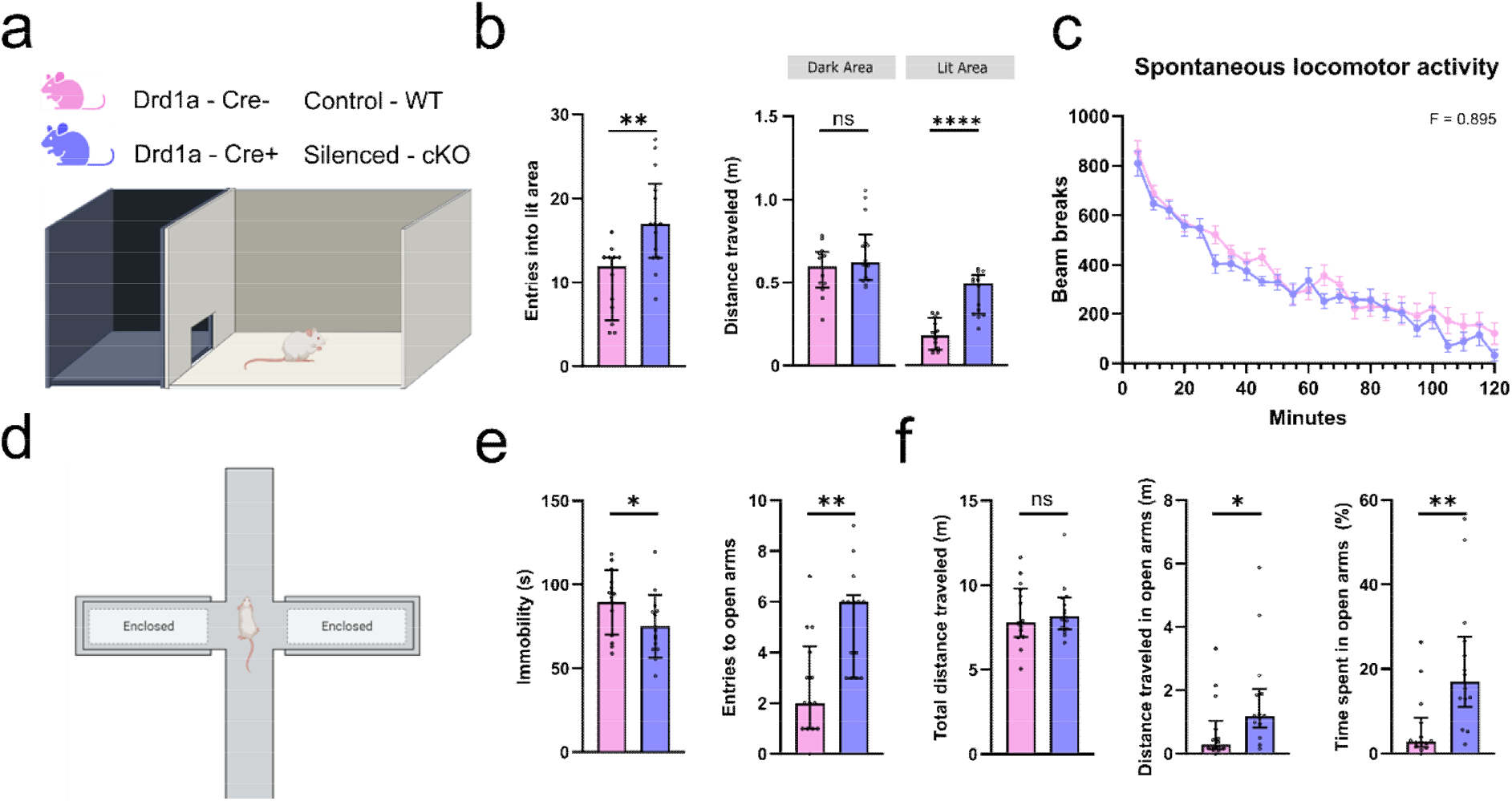
Anxious behaviour is altered after chronic manipulation of the cortical Drd1a-Cre+ neurons. **a)** Schematic of a Light-Dark Box (LDB) experimental protocol. On top, the colour coding for genotype of the control (Drd1a-Cre^-/-^:Ai14:Snap25^fl/fl^; WT) and Drd1a-Cre-silenced mice (Drd1a-Cre^+/-^:Ai14:Snap25^fl/fl^; cKO). **b)** cKO animals show an increase in entries into (n = 12; p = 0.0011;Mann Whitney test) and total movement inside the lit area (n = 12; p <0.0001 ; Mann Whitney test). **c)** Comparison of spontaneous locomotor activity in novel, photocell locomotor activity cages between WT and cKO animals shows no difference in the total movement of these animals **d)** Schematic set-up for the Elevated Plus Maze (EPM). **e)** cKO animals show a reduction in immobility after being placed in the EPM (n = 14; p = 0.0444; Mann Whitney test), as well as increased entries into the open arms (n = 14; p = 0.0016; Mann Whitney test). **f)** Total distance travelled was similar between genotypes, but both the distance moved in open arms (n = 14; p = 0.0141; Mann Whitney test), and the overall time spent in them (n = 14; p = 0.0012; Mann Whitney test) was significantly increased. All behavioural comparisons were tested by using Mann-Whitney nonparametric tests and all numbers were reported as the median. Error bars represent interquartile ranges, each datapoint is represented as a dot.

Both these results suggest a reduction in anxious behaviour following chronic silencing of Drd1a-Cre+ L6 neurons, which is more likely to reflect changes in higher association areas, including mPFC, rather than sensorimotor cortex. These Drd1a-Cre+ L6 neurons may thus provide a promising new target for the pharmacological treatment of anxiety disorders.

## Funding

MRC (ZM + AHZ), MRC (ZM, EM, FM), St John’s Research Centre Grant awarded to ZM, EM; Marie Skłodowska-Curie grant agreement No 890457-L6b (RT, EM, ZM); Dale fellowship from Wellcome Trust No 213465 (AL); Wellcome Trust Studentship (LG); Anatomical Society Studentship awarded to AH-S and ZM (KK);

## Acknowledgements

Dr Kristina Parley (lab manager ZM). Maxwell Biosystems (Zürich, Switzerland) for the high-density microelectrode array (MaxOne).

## Data availability

All data analysed in this study are available upon request from the corresponding authors (ZM & EM).

## Material and Methods

### Animals

All animal experiments were approved by a local ethical review committee and conducted in accordance with personal and project licenses under the UK Animals (Scientific Procedures) Act (1986). Mice were housed in a temperature-controlled room under a 12 h light/12 h dark cycle, with free access to food and water, and pups were kept with their dam until weaning age at P21, or the experimental endpoint if earlier. A total of 64 adult mice were used in this study. A mix of male and female animals (n=49 and n=15 respectively) were used in these experiments. Distinct Cre-recombinase-expressing strains were used for each experiment. The specifics of the genotypes used in each experiment are explained in detail in the corresponding section. A complete description of the animals used, and their genotype is shown in Table S1.

### Immunohistochemistry

Adult mice were anesthetised and culled by intraperitoneal pentobarbitone overdose (Animalcare, XVD135) followed by exsanguination and perfusion fixation using 0.1M phosphate buffered saline (PBS, pH=7.4) and 4% formaldehyde (PFA diluted in PBS, Sigma Aldrich, F8775). Brains were dissected, post-fixed for 24h in PFA, then stored in 0.1M PBS with 0.05% sodium azide at 4°C. Brains were embedded in 4.5% agarose, cut into 50μm coronal sections using a vibrating microtome (VT1000S, Leica Systems, Wetzlar, Germany), and placed into 24-well plates (one section/500μL/well). Tissues were washed 3×10min in 0.1MPBS, incubated in a blocking solution of 10% donkey serum (Sigma Aldrich, D9663) and 0.3% Triton™ X-100 (Thermo Scientific™, 85111) in 0.1M PBS (DS-10%;TX-0.3%;1XPBS) for 2hr at room temperature (RT), then with primary antibodies Rb α Cplx3 (1:1000, Synaptic Systems 122302) and Ms α Tbr1 (1:500, ProteinTech 66564-1-LG) in DS-5%;TX-0.3%;1XPBS for 48hr at 4°C. Sections were washed 3×10min, incubated with secondary antibodies Dk α Rb (488nm, 1:500, Invitrogen A21206) and Dk α Ms (647nm, 1:500, Invitrogen, A32787) in DS-5%;TX-0.3%;1XPBS for 2hr at RT, washed 3×10min in 0.1M PBS, incubated with 1:1000 4′,6-diamidino-2-phenylindole (DAPI) for 30min at RT, then washed 2×10min in 0.1M PBS. Tissues were mounted with 60μL ProLong™ Glass (Invitrogen™, P36984) per slide.

### Imaging and Analysis

Spinning-disk confocal microscopy (Olympus SpinSR SoRa) was conducted at 20X magnification, using cellSens (v4.1.1, Evident, Boston, MA, USA) to apply high-density focus maps with additional points as required. Images were pre-processed using a custom Fiji/ImageJ .ijm script (2.9.0 v1.54b). Coronal sections were imported with split channels at a downsampled factor of 4 (Series 3) followed by maximum-intensity z-stack projections, auto-brightness thresholding, and background subtraction. Outputs were imported into QuPath (v0.4.2) (Bankhead et al., 2017) where a custom Groovy script detected cells and exported two sets of images. The first contained .svg segmentations, which were re-formatted for Nutil compatibility (Yates et al., 2019) using another custom Fiji/ImageJ.ijm script (i.e., binary masks application, pixel inversion, and file type conversion to RGB .pngs). The second set of images were saved as RGB .pngs with 8 × downsampling for QuickNII compatibility (RRID:SCR_016854) (Puchades et al., 2019). Downsampled .pngs were formatted using QuickNII’s FileBuilder (Puchades et al., 2019) for subsequent section alignment and Allen atlas registration in QuickNII (2017 CCFv3) (Wang et al., 2020). Resulting .json anchoring files were imported to VisuAlign (v0.8 RRID:SCR_017978) for nonlinear refinement. Cell detections and VisuAlign outputs were imported into Nutil (Groeneboom et al., 2020).Brain regions of interest were defined using a custom Excel sheet. Object reports were generated with a minimum size of 4 pixels, point cloud density of 4, without object splitting, and extracting all coordinates. A custom MATLAB script (R2022b, MathWorks, Natick, MA, USA) re-formatted Nutil .csv outputs for statistical analysis and graphical representation in GraphPad Prism (v.9.3.1, San Diego, CA, USA).

### *In vitro* patch-clamp electrophysiology

*In vitro* electrophysiological experiments were done on a genetically modified strain (Tg(Drd1a-Cre)FK164Gsat/Mmucd), that selectively expresses Cre-recombinase in a subpopulation of layer 6 neurons across the entire cortical mantle. This L6-specific Cre line is crossed with a td-Tomato reporter line (Ai14). 18 adult mice aged 22.83 ± 6.03 were used for single-cell patch-clamp electrophysiological recordings (for details see Supplementary Table 1). An average of two L6 neurons were recorded per animal. Animals were anesthetized using 4% isoflurane followed by decapitation, and the brains were extracted in cold sucrose solution (40 mM NaCl, 3 mM KCl, 7.4 mM MgSO_4_.7H_2_O, 150 mM sucrose, 1 mM CaCl_2_, 1.25 mM NaH_2_PO_4_, 25 mM NaHCO_3_, and 15 mM glucose; osmolality 300 ± 10 mOsmol/kg).

Coronal cortical slices (250 - 300 μm thick) were cut using a vibratome (Leica VT1200S) and placed in an interface chamber containing artificial cerebrospinal fluid (aCSF) (126 mM NaCl, 3.5 mM KCl, 2 mM MgSO_4_.7H_2_O, 1.25 mM NaH_2_PO_4_, 24 mM NaHCO_3_, 2 mM CaCl_2_, and 10 mM glucose; osmolality 300 ± 10 mOsmol/kg) for the duration of the experiment. All solutions were bubbled with carbogen gas (95% O2/ 5% CO2). Whole-cell current-clamp recordings of mPFC neurons were performed in a submerged chamber (∼ 32 °C) using borosilicate glass pipettes (5–12 MΩ) filled with internal solution (110 mM K-gluconate, 40 mM HEPES, 2 mM ATP-Mg, 0.3 mM GTP-NaCl, 4 mM NaCl, 3–4 mg/ml biocytin; pH ∼7.2; osmolality 270–290 mOsmol/kg). Electrical signals were amplified with a Multiclamp 700 B amplifier (Molecular Devices, Foster City, CA), low-pass filtered at 10 kHz and digitized at 20 kHz using a Digidata 1440A (Molecular Devices). Hyperpolarizing and depolarizing square current pulses were applied in order to quantify intrinsic properties of the recorded neuron, with these step protocols performed at baseline and 2, 5 and 10 minutes after application of 200 nM YNT-185. The recordings were extracted and processed using Igor Pro 6 (Wavemetrics) and later analysed with custom-made Python scripts. All statistics were done using GraphPad Prism 10.

### *In vivo* injections

*In vivo* Orexin-B injections were performed on the same genetically modified strain of *Tg(Drd1a-Cre)FK164Gsat/Mmucd* used in the patch-clamp experiments. A total of 9 animals were used, aged 78.80 ± 31.71 days. The mice were anaethetised for the duration of the experiment using 2% isoflurane/medical oxygen mixture, implanted with a 16-channel linear electrode (Neuronexus) targeting mPFC (1.94 mm antero-posterior, 0.4 mm lateral and 1.75 mm dorso-ventral), with extracellular voltage signals acquired at 30 kHz using an RHD2164 64-channel amplifier board connected to an RHD2000 USB interface board and controlled by RHX Data Acquisition software (Intan Technologies). The animals were injected with 1-2 µL of either a 0.345 mM Orexin-B solution or a 1-2 µL saline solution in the left lateral ventricle (−0.1 mm antero-posterior, 0.75 mm lateral and: 1.90 mm dorso-ventral). A 15-minute recording before, and a 20-minute recording after the injection were performed in all animals. The first 5 minutes of the recording following the Orexin-B administration were not used for the analysis. Recordings were extracted and processed with custom-made Python scripts and all statistics were calculated with GraphPad Prism 10.

### Multielectrode electrophysiology

The network effect of silencing the L6-Drd1a population was evaluated in a Drd1a-Cre^+/-^ :Ai14:Snap25^fl/fl^ genetically modified strain, and compared with Cre negative controls Drd1a-Cre^-/-^ :Ai14:Snap25^fl/fl^. These Drd1a-Cre^+/-^:Ai14:Snap25^fl/fl^ animals express Td-Tomato and have Snap25 ablated in the Drd1a positive neurons, leading to severe reductions in evoked synaptic vesicle release from these neurons (Hoerder-Suabedissen et al., 2019). As an additional control, a similar genetical modification was performed in Rbp4-Cre:Ai14:Snap25^fl/fl^ which mirrors the Snap25 ablation of the previous population, but in Rbp4-Cre positive neurons (Kozorovitskiy et al., 2012).

9 adult mice aged 76.14 ± 5.03 were used. Brains from the animals were retrieved and the tissue was sliced using the same protocol as the one described for the *in vitro* whole-cell current-clamp experiments. An average of 3 slices per animal were used for each group. All experiments were performed by experimenters blind to the genotypes of the animals. Recordings were done using a high-density microelectrode array (MaxOne) from Maxwell Biosystems (Zürich, Switzerland). The MEA provides 26,400 electrodes in a 3.85 × 2.10 mm^2^ large sensing area with an electrode pitch of 17.5 μm (Müller et al., 2015). A combination of up to 1,024 electrodes can be recorded from simultaneously, with a sampling rate of 20 kHz. All recordings were 5 minutes long, each condition was done by triplicate. Baseline recordings were done using a continuous administration of 200 µM 4-AP in aCSF for 20 minutes before recordings started. Afterwards, the solution was changed for a 200 nM YNT-185 and 200 µM 4-AP in aCSF, following the same protocol as before. The recordings were extracted and pre-processed using custom-made Matlab and Python scripts. All statistics were done using GraphPad Prism 10.

### Behavioural experiments

The behavioural evaluation was performed using the same genetically modified strain as used for the high-density multielectrode electrophysiology: Drd1a-Cre^+/-^:Ai14:Snap25^fl/fl^ versus Drd1a-Cre^-/-^ :Ai14:Snap25^fl/fl^. Anxiety-related behaviour was assessed using two protocols: i) the elevated plus maze (EPM), which consisted of an elevated platform at 50 cm from the floor and composed of four arms 30 cm long and 5 cm wide, and a 5 by 5 cm central zone. The arms were divided into two pairs, one pair of them enclosed by a 30 cm lateral wall and the other pair open. For this test, animals were placed in the central zone facing one of the open arms and allowed to explore the arms freely. Time spent in open arms and number of entries into open arms were recorded for 5 minutes using AnyMaze tracking system (Stoelting). ii) the light-Dark box (LDB), which consisted of a 44 by 21 by 21 cm cage divided into light and dark chambers by a 13 cm long and 5 cm high opening. One area was made dark with black spray paint, covered from the external environment, and maintained without illumination. The other compartment was open and brightly lit. The lit area corresponded to 2/3 of the total area of the box. Measurement of spontaneous locomotor activity (LMA) in a novel environment was conducted in plexiglass cages (20 × 35 cm), with movement detected with an infrared photobeams system (PAS-Open Field, San Diego Instruments) over the course of 2 h every minute. All behavioural experiments were performed by experimenters blind to the genotypes of the animals.

## Notes

### Competing Interest Statement

The authors have declared no competing interest.

## Reference

Bankhead, P., Loughrey, M. B., Fernandez, J. A., Dombrowski, Y., McArt, D. G., Dunne, P. D., McQuaid, S., Gray, R. T., Murray, L. J., Coleman, H. G., James, J. A., Salto-Tellez, M., & Hamilton, P. W. (2017). QuPath: Open source software for digital pathology image analysis. Sci Rep, 7(1), 16878. 10.1038/s41598-017-17204-5

Belzung, C., & Griebel, G. (2001). Measuring normal and pathological anxiety-like behaviour in mice: a review. Behavioural Brain Research, 125, 141–149.

Bloem, B., Poorthuis, R. B., & Mansvelder, H. D. (2014). Cholinergic modulation of the medial prefrontal cortex: the role of nicotinic receptors in attention and regulation of neuronal activity. Front Neural Circuits, 8, 17. 10.3389/fncir.2014.00017

Bourassa, J., Pinault, D., & Deschenes, M. (1995). Corticothalamic projections from the cortical barrel field to the somatosensory thalamus in rats: a single-fibre study using biocytin as an anterograde tracer. Eur J Neurosci, 7(1), 19–30. 10.1111/j.1460-9568.1995.tb01016.x

Bourin, M., & Hascoet, M. (2003). The mouse light/dark box test. Eur J Pharmacol, 463(1-3), 55–65. 10.1016/s0014-2999(03)01274-3

Clancy, B., & Cauller, L. J. (1999). Widespread projections from subgriseal neurons (layer VII) to layer I in adult rat cortex. The Journal of Comparative Neurology, 407(2), 275–286. 10.1002/(sici)1096-9861(19990503)407:2<275::Aid-cne8>3.0.Co;2-0

Cole, S., Keefer, S. E., Anderson, L. C., & Petrovich, G. D. (2020). Medial Prefrontal Cortex Neural Plasticity, Orexin Receptor 1 Signaling, and Connectivity with the Lateral Hypothalamus Are Necessary in Cue-Potentiated Feeding. J Neurosci, 40(8), 1744–1755. 10.1523/JNEUROSCI.1803-19.2020

Crawley, J., & Goodwin, F. K. (1980). Preliminary Report of a Simple Animal Behavior Model for the Anxiolytic Effects of Benzodiazepines. Pharmacology Biochemistry & Behavior, Vol. 13, 167–170.

Geiger, J. R. P., & Jonas, P. (2000). Dynamic Control of Presynaptic Ca2 Inflowby Fast-Inactivating K Channels in Hippocampal Mossy Fiber Boutons. Neuron, 28, 927–939.

Groeneboom, N. E., Yates, S. C., Puchades, M. A., & Bjaalie, J. G. (2020). Nutil: A Pre- and Post-processing Toolbox for Histological Rodent Brain Section Images. Front Neuroinform, 14, 37. 10.3389/fninf.2020.00037

Henning, T., Bandiera, S., & Molnár, Z. (2023). Towards the Transcriptionally based Classification of L6b in the Adult Mouse Brain. In Neocortical Neurogenesis in Development and Evolution (pp. 317–330). 10.1002/9781119860914.ch16

Hoerder-Suabedissen, A., Hayashi, S., Upton, L., Nolan, Z., Casas-Torremocha, D., Grant, E., Viswanathan, S., Kanold, P. O., Clasca, F., Kim, Y., & Molnar, Z. (2018). Subset of Cortical Layer 6b Neurons Selectively Innervates Higher Order Thalamic Nuclei in Mice. Cereb Cortex, 28(5), 1882–1897. 10.1093/cercor/bhy036

Hoerder-Suabedissen, A., Korrell, K. V., Hayashi, S., Jeans, A., Ramirez, D. M. O., Grant, E., Christian, H. C., Kavalali, E. T., Wilson, M. C., & Molnar, Z. (2019). Cell-Specific Loss of SNAP25 from Cortical Projection Neurons Allows Normal Development but Causes Subsequent Neurodegeneration. Cereb Cortex, 29(5), 2148–2159. 10.1093/cercor/bhy127

Karimi, S., Hamidi, G., Fatahi, Z., & Haghparast, A. (2019). Orexin 1 receptors in the anterior cingulate and orbitofrontal cortex regulate cost and benefit decision-making. Prog Neuropsychopharmacol Biol Psychiatry, 89, 227–235. 10.1016/j.pnpbp.2018.09.006

Katzman, M. A., & Katzman, M. P. (2022). Neurobiology of the Orexin System and Its Potential Role in the Regulation of Hedonic Tone. Brain Sci, 12(2). 10.3390/brainsci12020150

Killackey, H. P., & Sherman, S. M. (2003). Corticothalamic Projections from the Rat Primary Somatosensory Cortex. The Journal of Neuroscience, 23, 7381–7384.

Kole, M. H., Letzkus, J. J., & Stuart, G. J. (2007). Axon initial segment Kv1 channels control axonal action potential waveform and synaptic efficacy. Neuron, 55(4), 633–647. 10.1016/j.neuron.2007.07.031

Kozorovitskiy, Y., Saunders, A., Johnson, C. A., Lowell, B. B., & Sabatini, B. L. (2012). Recurrent network activity drives striatal synaptogenesis. Nature, 485(7400), 646–650. 10.1038/nature11052

Kress, G. J., & Mennerick, S. (2009). Action potential initiation and propagation: upstream influences on neurotransmission. Neuroscience, 158(1), 211–222. 10.1016/j.neuroscience.2008.03.021

Laviolette, S. R., Lipski, W. J., & Grace, A. A. (2005). A subpopulation of neurons in the medial prefrontal cortex encodes emotional learning with burst and frequency codes through a dopamine D4 receptor-dependent basolateral amygdala input. J Neurosci, 25(26), 6066–6075. 10.1523/JNEUROSCI.1168-05.2005

Lodge, D. J. (2011). The medial prefrontal and orbitofrontal cortices differentially regulate dopamine system function. Neuropsychopharmacology, 36(6), 1227–1236. 10.1038/npp.2011.7

Müller, J., Ballini, M., Livi, P., Chen, Y., Radivojevic, M., Shadmani, A., Viswam, V., Jones, I. L., Fiscella, M., Diggelmann, R., Stettler, A., Frey, U., Bakkum, D. J., & Hierlemann, A. (2015). High-resolution CMOS MEA platform to study neurons at subcellular, cellular, and network levels. Lab Chip, 15(13), 2767–2780. 10.1039/c5lc00133a

Puchades, M. A., Csucs, G., Ledergerber, D., Leergaard, T. B., & Bjaalie, J. G. (2019). Spatial registration of serial microscopic brain images to three-dimensional reference atlases with the QuickNII tool. PLoS One, 14(5), e0216796. 10.1371/journal.pone.0216796

Shu, Y., Yu, Y., Yang, J., & McCormick, D. A. (2007). Selective control of cortical axonal spikes by a slowly inactivating K current. PNAS, 104(27).

Thomson, A. M. (2010). Neocortical layer 6, a review. Front Neuroanat, 4, 13. 10.3389/fnana.2010.00013

Viswanathan, S., Sheikh, A., Looger, L. L., & Kanold, P. O. (2017). Molecularly Defined Subplate Neurons Project Both to Thalamocortical Recipient Layers and Thalamus. Cereb Cortex, 27(10), 4759–4768. 10.1093/cercor/bhw271

Walf, A. A., & Frye, C. A. (2007). The use of the elevated plus maze as an assay of anxiety-related behavior in rodents. Nat Protoc, 2(2), 322–328. 10.1038/nprot.2007.44

Wang, Q., Ding, S. L., Li, Y., Royall, J., Feng, D., Lesnar, P., Graddis, N., Naeemi, M., Facer, B., Ho, A., Dolbeare, T., Blanchard, B., Dee, N., Wakeman, W., Hirokawa, K. E., Szafer, A., Sunkin, S. M., Oh, S. W., Bernard, A., … Ng, L. (2020). The Allen Mouse Brain Common Coordinate Framework: A 3D Reference Atlas. Cell, 181(4), 936–953 e920. 10.1016/j.cell.2020.04.007

Wang, Z., & Mengoni, P. (2022). Seizure classification with selected frequency bands and EEG montages: a Natural Language Processing approach. Brain Inform, 9(1), 11. 10.1186/s40708-022-00159-3

Yates, S. C., Groeneboom, N. E., Coello, C., Lichtenthaler, S. F., Kuhn, P. H., Demuth, H. U., Hartlage-Rubsamen, M., Rossner, S., Leergaard, T., Kreshuk, A., Puchades, M. A., & Bjaalie, J. G. (2019). QUINT: Workflow for Quantification and Spatial Analysis of Features in Histological Images From Rodent Brain. Front Neuroinform, 13, 75. 10.3389/fninf.2019.00075

Zolnik, T. A., Bronec, A., Ross, A., Staab, M., Sachdev, R. N. S., Molnar, Z., Eickholt, B. J., & Larkum, M. E. (2023). Layer 6b controls brain state via apical dendrites and the higher-order thalamocortical system. Neuron. 10.1016/j.neuron.2023.11.021

Zolnik, T. A., Ledderose, J., Toumazou, M., Trimbuch, T., Oram, T., Rosenmund, C., Eickholt, B. J., Sachdev, R. N. S., & Larkum, M. E. (2020). Layer 6b Is Driven by Intracortical Long-Range Projection Neurons. Cell Rep, 30(10), 3492–3505 e3495. 10.1016/j.celrep.2020.02.044

