## Supplementary figures 1-5 for "An orexin-sensitive subpopulation of layer 6 neurons regulates cortical excitability and anxiety behaviour"

### Supplementary Information

#### Figure S1: Determination L6a & L6b

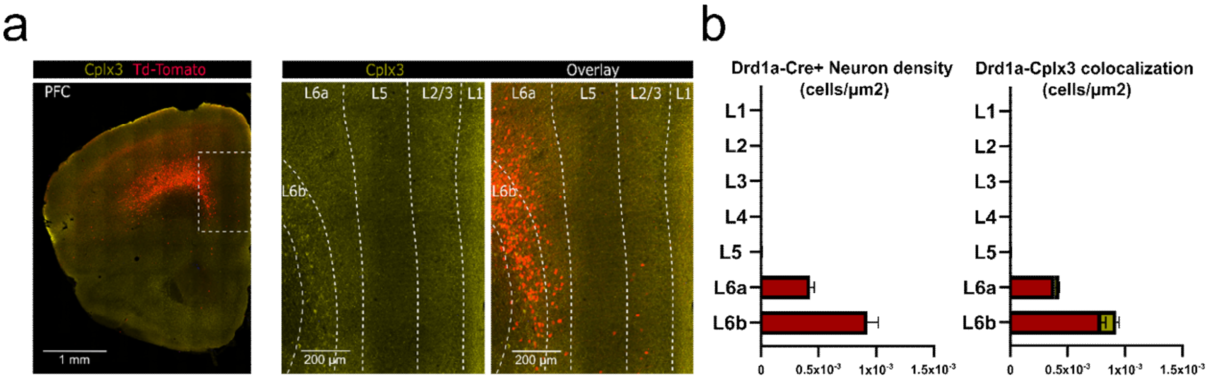


**a)** Representative image of Cplx-3 immunostained sections of Drd1a-Cre:Ai14 mice. These animals express Td-Tomato in the Drd1a-Cre+ neurons. Determination of the L6a-L6b border was done using Cplx3 staining (yellow). **b)** There is a higher density of Drd1a-Cre+ neurons in L6b, although is not exclusive to this layer. The proportion of Drd1a+/Cplx3+ neurons is higher in the deeper parts of L6.

#### Figure S2: Spiking activity in Drd1a-Cre+ and Drd1a-Cre- neurons in PFC layer 6

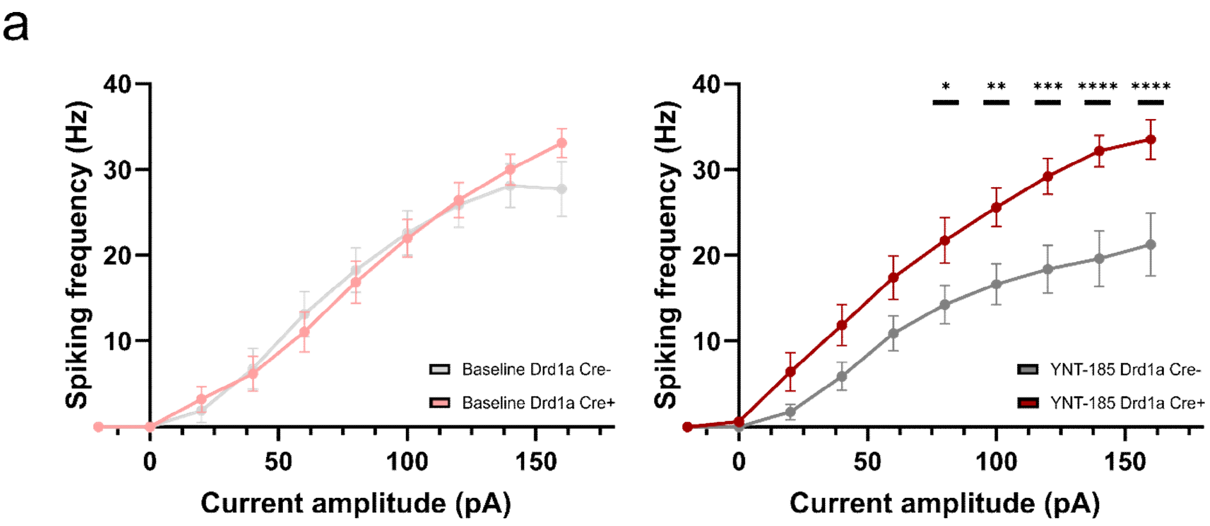


**a)** The input/output curve shows the changes in spiking activity after increasing stimulation during baseline (aCSF) (Left) and after administration of orexin agonists (YNT-185) (Right). Baseline activity shows no difference between distinct genotypes. Drd1a-Cre+ neurons respond significantly more to the administration of YNT-185 as compared to the Drd1a-Cre- neurons ($n = 15$; $p < 0.0001$; Mixed effect analysis with Šídák's multiple comparisons). All numbers were reported as the mean, error bars represent SEM.

### Figure S3: Action potential properties of recorded Drd1a-Cre+ neurons

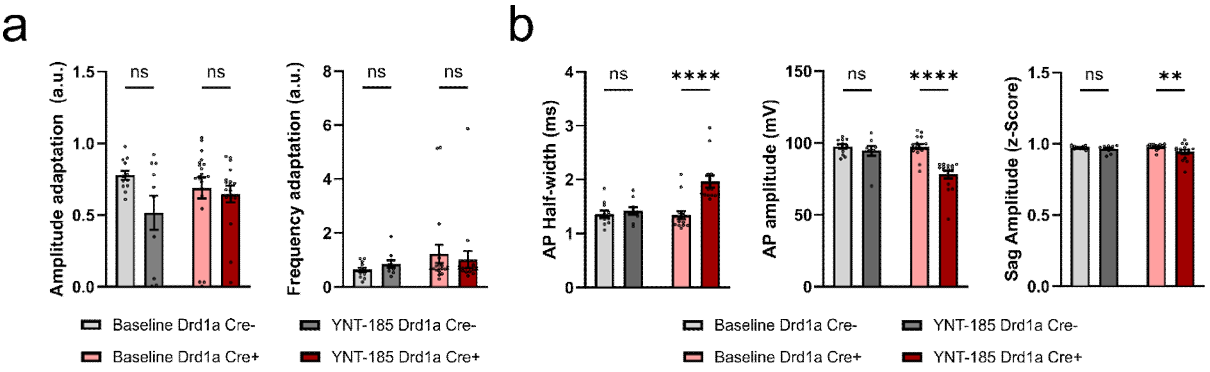


**a)** Administration of YNT-185 had no significant effect on the evoked action potentials' frequency or amplitude adaptation. We also observed no difference between Drd1a-Cre+ and Drd1a-Cre- neurons. **b)** Consistent with previous reports of frequency-dependent changes, action potentials showed a significant increase in the half-width and a decrease in the spike amplitude (n = 15; p <0.0001; Mixed effect analysis with Šídák's multiple comparisons). All numbers were reported as the mean, error bars represent SEM.

### Figure S4: Spiking frequency

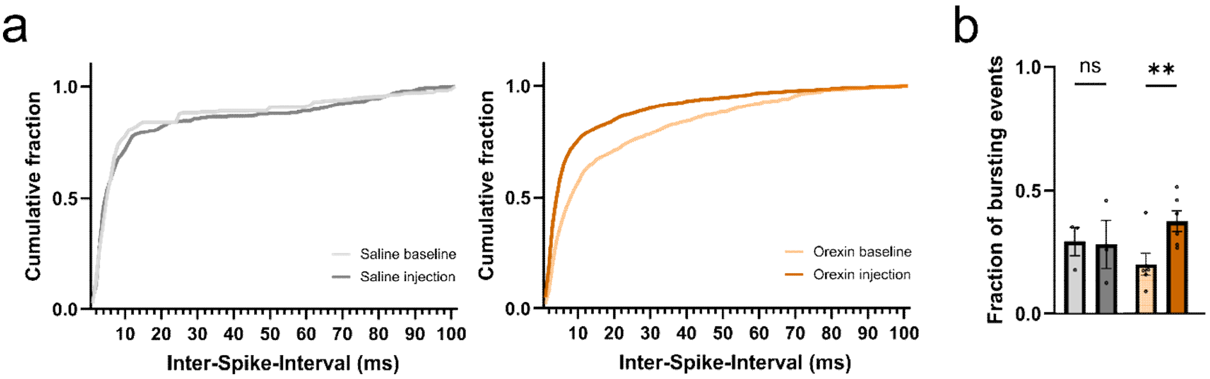


**a)** Orexin-B intraventricular injections increase the number of evoked bursting events. Compared to a saline control (Left), Orexin-B injections increased the proportion of events with a inter-spike-interval smaller than 50 ms. **b)** The increase in the number of bursting events was significant after the orexin-B injection The increase in evoked spiking activity was not reflected as an increment in bursting activity for any of the injections (n = 6; p = 0.0088; Mixed effect analysis with Šídák's multiple comparisons). All numbers were reported as the mean, error bars represent SEM.

**Figure S5: Multielectrode activation of L5 silenced animals**

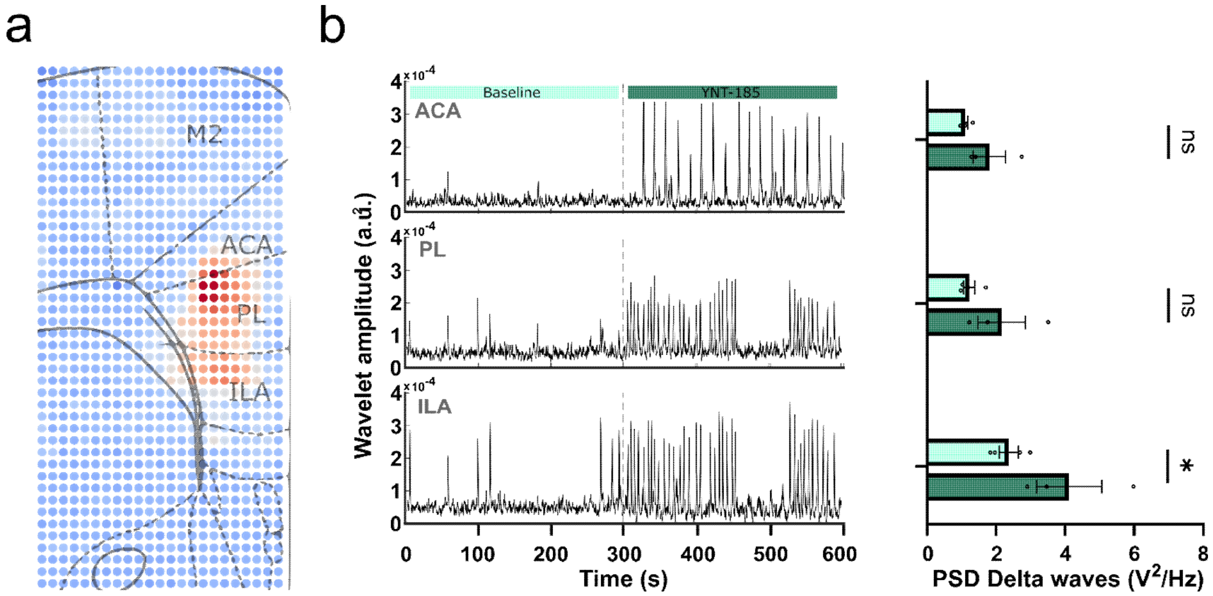


**a)** PSD analysis of the network response to Orexin agonist in the layer 5 silenced animals (Rbp4-Cre+/+:Snap25fl/fl). The administration of orexin significantly increases the delta activity in mPFC. The silencing of layer 5 had no effect in the orexinergic activation and, together with the findings of Figure 4, demonstrates that the orexinergic activation of the neocortex depends on the activation of layer 6 Drd1a-Cre+ neurons. **b)** Wavelet transform of the recordings shows an increase in activity across time in all mPFC areas after YNT-185 administration. All comparisons were done using Mixed-effect Two-Way ANOVA with Šidák correction. All numbers were reported as the mean with SEM. Error bars represent SEM, each datapoint is represented as a dot
